# Biological misinterpretation of transcriptional signatures in tumour samples can unknowingly undermine mechanistic understanding and faithful alignment with preclinical data

**DOI:** 10.1101/2022.04.15.488354

**Authors:** Natalie C Fisher, Ryan M Byrne, Holly Leslie, Colin Wood, Assya Legrini, Andrew J Cameron, Baharak Ahmaderaghi, Shania Corry, Sudhir Malla, Raheleh Amirkhah, Aoife McCooey, Emily Rogan, Keara L Redmond, Svetlana Sakhnevych, Enric Domingo, James Jackson, Maurice B Loughrey, Simon Leedham, Tim Maughan, Mark Lawler, Owen J Sansom, Felicity Lamrock, Viktor H Koelzer, Nigel Jamieson, Philip D Dunne

## Abstract

Precise mechanism-based gene expression signatures (GESs) have been developed in appropriate *in vitro* and *in vivo* model systems, to identify important cancer-related signalling processes. However, some GESs originally developed to represent specific disease processes, primarily with an epithelial cell focus, are being applied to heterogeneous tumour samples where the expression of the genes in the signature may no longer be epithelial-specific. Therefore, unknowingly, even small changes in tumour stroma percentage can directly influence GESs, undermining the intended mechanistic signalling.

Using colorectal cancer as an exemplar, we deployed numerous orthogonal profiling methodologies, including laser capture microdissection, flow cytometry, bulk and multiregional biopsy clinical samples, single cell RNAseq and finally spatial transcriptomics, to perform a comprehensive assessment of the potential for the most widely-used GESs to be influenced, or confounded, by stromal content in tumour tissue. To complement this work, we generated a freely-available resource, ConfoundR; https://confoundr.qub.ac.uk/, that enables users to test the extent of stromal influence on an unlimited number of the genes/signatures simultaneously across colorectal, breast, pancreatic, ovarian and prostate cancer datasets.

Findings presented here demonstrate the clear potential for misinterpretation of the meaning of GESs, due to widespread stromal influences, which in-turn can undermine faithful alignment between clinical samples and preclinical data/models, particularly cell lines and organoids, or tumour models not fully recapitulating the stromal and immune microenvironment. As such, efforts to faithfully align preclinical models of disease using phenotypically-designed GESs must ensure that the signatures themselves remain representative of the same biology when applied to clinical samples.

## Introduction

Although the publication of gene expression-based signatures (GES) continues to grow each year in the research setting, these published signatures rarely make any clinical impact.^1^ In addition to potentially-addressable technical confounders, such as sample size issues or lack of validation cohorts, the biology underpinning the signature may also expose a critical weakness in current translational bioinformatics research pipelines, when applied to clinical samples either in retrospective collections or prospective trials. This is particularly pertinent as researchers now have unparalleled access to cancer datasets that can be routinely characterised using the tens of thousands of GESs already available in molecular databases such as TCGA.^2^ We have previously demonstrated the confounding effects of stroma on molecular subtypes in colorectal cancer,^3,4^ alongside specific influences of the tumour microenvironment on EMT-related signatures.^5,6^

A number of recent studies have highlighted the characterisation required to ensure faithful alignment between human tumours and preclinical models, in terms of the biological signalling and therapeutic responses in both.^7-9^ Integrity and robustness in aligning models with human tumours is critical in the era of precision medicine, where treatments are tailored for the biology underpinning specific cancer subtypes. Furthermore, signature development and testing is increasingly performed in disease-matched pre-clinical models, using *in vitro, in vivo* or *ex vivo* systems, enabling almost absolute control over the experimental conditions employed during biology-driven GES development.^10-12^ Although such “clean” models are exquisitely suited for precise identification and characterisation of discrete mechanistic signalling, when compared to the relative unpredictable nature of diagnostic sample acquisition, differences in the epithelial, immune and stromal composition between the models and clinical samples^13^ has the potential to confound our understanding and interpretation of these signatures in specific domains. While this will in no way alter the prognostic/predictive statistical value of such signatures, differences in cellular composition and tumour stroma percentage (TSP) are rarely accounted for during the interpretation of the true biological meaning of the GES result in bulk tumour datasets. Conversely, when biomarkers of prognosis, response or molecular subtypes are identified from tumour datasets, approaches to reverse-translate these findings into pre-clinical models introduces the potential for assessment of these genes/signatures in lineages that do not represent the true cellular source of the signalling in clinical samples. Again, it is important to note that the *statistical correlation* between a specific biomarker/signature and a clinical variable like relapse, are in no way weakened if the end-user does not accurately consider the true *biological interpretation* of the signature itself. While biological researchers understand that correlation does not always equate to causation,^14^ there remains a potential gap in our understanding when interpretation of GESs can be influenced by the cellular composition of a tumour sample. The potential for misinterpretation is an issue that has become even more important in the precision medicine era,^15^ where it is now essential that therapeutic targeting is based on robust and accurate mechanistic-driven evidence. In order to successfully translate pre-clinical efficacies into clinical benefit, testing of therapeutics must be performed in models that are representative of specific patient subtypes.^7^

To examine if variations in TSP can distort GES results, and in-turn lead to biological misinterpretation, we performed a comprehensive assessment and quantification of the extent that stromal composition in bulk tumours can skew the expression levels of n=7835 of commonly employed gene sets and signatures in cancer research. Using a combination of discovery and independent validation cohorts, including tissues from laser capture microdissection, flow cytometry, bulk clinical samples, single cell RNAseq and finally spatial transcriptomics, enabled a detailed interrogation of widely-used transcriptomic signatures to enumerate the extent to which stromal composition can confound their classification. The pan-cancer nature of these findings were subsequently assessed across a number of publicly-available laser capture microdissection datasets derived from pancreatic, breast, ovarian and prostate cancer. Furthermore, to ensure that our findings can be widely applied, we have developed the freely-available ConfoundR on-line resource; https://confoundr.qub.ac.uk/, which gives a user the ability to quickly and easily interrogate the potential confounding effects on any individual gene, combination of genes, and GES across colorectal, pancreatic, breast, ovarian and prostate cancer datasets.

## Results

### Initial characterisation of tumour epithelium and stromal datasets

To assess the influence that TSP has on commonly used transcriptional signatures, we designed a study to identify, characterise, and orthogonally validate the tumour microenvironment (TME) compartments and lineages associated with specific transcriptional signatures within primary colorectal cancer (CRC). This approach utilised a series of independent primary tumour samples that had undergone laser capture microdissection (LCM), to segregate tissue into epithelial and stromal components, for discovery (n=26 samples from n=13 tumours; GSE35602) and validation (n=16 samples from n=8 tumours; GSE31279) (Figure 1A). Further delineation of cell-type-specific transcriptional signalling was performed using transcriptional data generated from FACS-purified epithelial, fibroblast, endothelial and leukocyte cell populations from CRC resections (n=6 tumours, n=24 populations; GSE39396) (Figure 1A).

**Figure 1.**
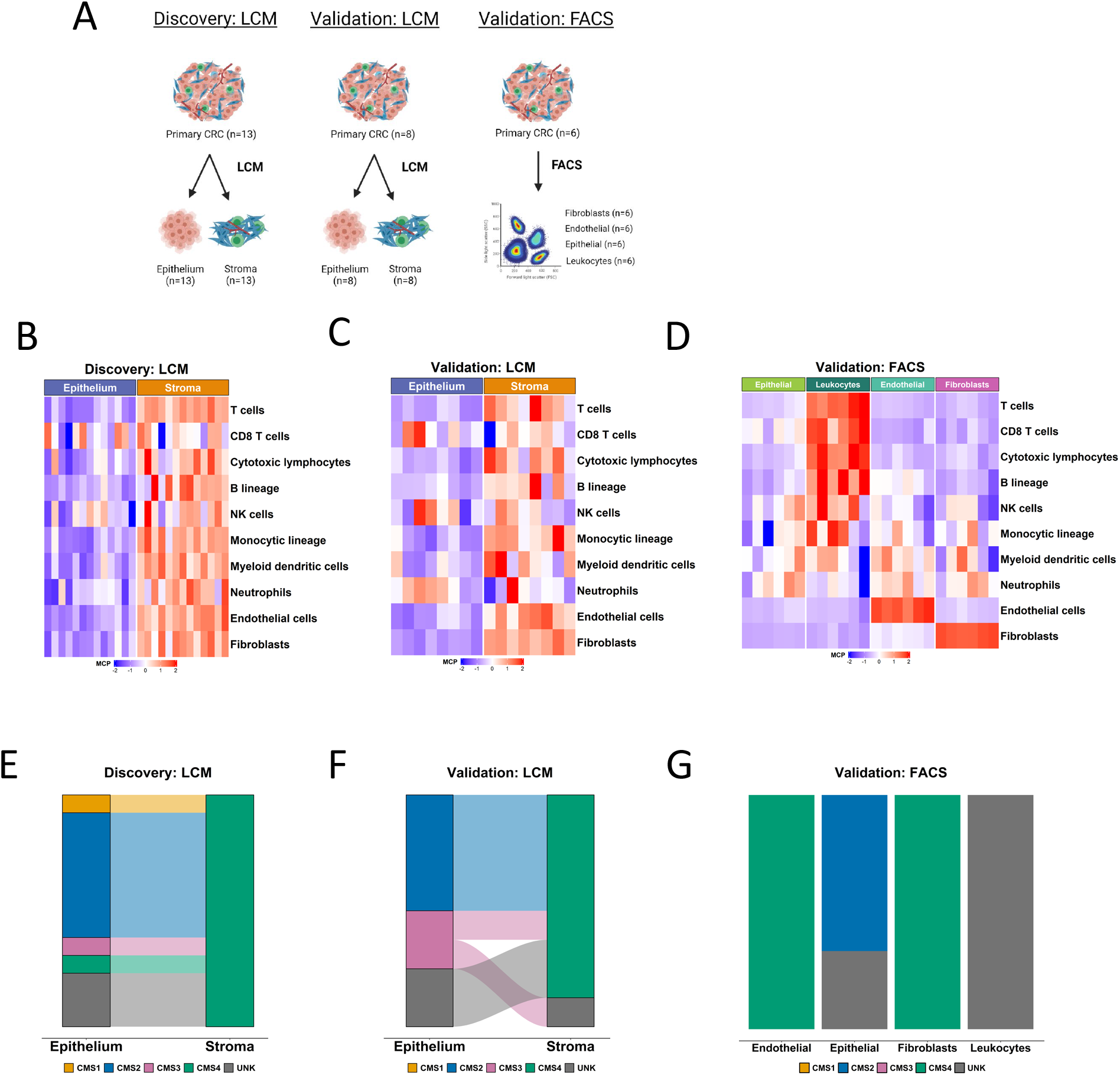
Initial characterisation of tumour epithelium and stromal datasets. **A** Schematic of the segregation strategies in the discovery and validation cohorts, drawn using BioRender. **B** Heatmap of MCP-counter scores for the laser capture microdissected (LCM) discovery cohort, according to epithelium and stromal regions. **C** Heatmap of MCP-counter scores for the LCM validation cohort, according to epithelium and stromal regions. **D** Heatmap of MCP-counter scores for the FACS validation cohort. **E** CMS classifications (using CMSclassifier) for the matched epithelium and stroma samples in the laser capture microdissected discovery cohort. **F** CMS calls (using CMSclassifier) for the matched epithelium and stroma samples in the laser capture microdissected validation cohort. **G** CMS calls (using CMSclassifier) for the four lineages in the FACS validation cohort.

To confirm the purity of these datasets, we utilised the microenvironment cell population (MCP)-counter algorithm^16^ to assign single sample scores for n=10 stromal (fibroblasts, endothelial cells) and immune lineages (T cells, CD8 T cells, cytotoxic lymphocytes, B lineages, natural killer (NK) cells, monocytic lineages, myeloid dendritic cells, neutrophils) to each individual sample (Figure 1B-D). In the LCM cohorts, these analyses confirmed that the majority of TME lineage signatures are exclusively stromal, particularly those aligned to fibroblast and endothelial cells (Figure 1B, C). Although most immune lineages appeared to align to the stroma, we did observe signalling indicative of CD8 T cells, NK cells and neutrophils within the epithelial compartment, indicative of intra-epithelial infiltration of these specific immune lineages (Figure 1B, C). In line with this LCM data, within the FACS cohort we observed fibroblast and endothelial populations aligned exclusively to the MCP-counter signature for fibroblasts and endothelial cells respectively, supporting the suitability of our approach and the utility of the MCP-counter signatures (Figure 1D). While T cell, CD8 T cell, cytotoxic lymphocyte and B cell lineage scores all closely aligned to the purified leukocyte population as expected, we did observe signalling indicative of NK cells, myeloid dendritic cells and neutrophils in non-leukocyte populations, suggesting that there was some crossover in these specific populations during cell sorting for epithelial cells (EPCAM+), leukocytes (CD45+), fibroblasts (FAP+) and endothelial cells (CD31+), or that the signatures cannot be used for precise enumeration of these lineages in CRC tissue.

### Association of CRC molecular subtypes with stromal components

A number of studies including our own have identified the stromal and immune contributions to the CRC consensus molecular subtypes (CMS)^3,4,17,18^, in particular to CMS1 and CMS4, however the relative contributions of TME compartments and specific lineage contributing to CMS calls using the original classifier have not been detailed. To test this, we classified the epithelial and stromal components from each tumour using the CMSclassifier^17^ algorithm, where we found that with the exception of one unclassified samples (UNK), the stroma was exclusively classified as CMS4 in the LCM cohorts (Figure 1E, F), as were both the purified fibroblast and endothelial lineages in the FACS cohort (Figure 1G), suggesting that transcriptional signalling from these components alone, even in the absence of the epithelial transcriptome, is sufficient for tumour classification as CMS4, the group with the worst prognosis in CRC.

When the epithelium was examined, with the exception of two samples, we observed a strong tendency for classification of CMS2 and CMS3, both well-characterised epithelial-rich subtypes, across the LCM and FACS cohorts (Figure 1E, F, G). In contrast to the association between stromal/endothelial cells and CMS4, when the leukocyte FACS purified population calls were assessed, we observed uniform unknown/unclassified assignments, indicating that the presence of immune infiltration alone is not sufficient for classification of a tumour as an immune-rich CMS1 tumour (Figure 1G) and more complex histological features involving tumour infiltrating lymphocytes and epithelial components are required. Furthermore, these issues remain apparent when using the CMScaller classifier,^19^ specifically modified to classify epithelial-based pre-clinical models according to CMS (Supplementary Figure 1).

### Stromal influence on widely used transcriptional signatures

While individual studies have highlighted the stromal origins of a number of key genes/proteins, using methods similar to MCP, it remains unknown how influential the stromal transcriptome is on some of the most widely-employed GESs. To investigate this, we performed pair-wise gene set enrichment analysis (GSEA)^20^ comparing epithelium to stroma using one of the most commonly used pathway/ontology collections, the Molecular Signatures Database (MSigDB)^21^ of n=50 “Hallmarks” (Supplementary Figure 2A, B). By performing these analyses in both LCM cohorts in tandem, we observed that n=21 Hallmarks were significantly (p_adj_ <0.02; more stringent that the accepted 0.25 cut-off) and consistently associated with either stroma (n=17) or epithelium (n=4) across both LCM cohorts (Figure 2A). These findings were further validated using single-sample GSEA (ssGSEA) in the FACS cohort (Supplementary Figure 2C), where the n=17 stromal-associated and n=4 epithelial-associated Hallmarks were again enriched in the corresponding cell populations (Figure 2B). Despite being developed and named to classify samples associated with specific biology, these analyses reveal the signalling underpinning of these signatures may be entirely, albeit unintentionally, misinterpreted due to the confounding effects of the stromal transcriptome in bulk tumour data. To ensure that this confounding effect is not an artefact of the Hallmark signatures specifically, we performed the same analyses using the n=186 KEGG and n=7481 gene ontology biological processes (GO BP) signatures, where again we found widespread stromal influence in 50/186 (Supplementary Figure 2D) and 949/7481 (Supplementary Table 1) signatures consistently in both cohorts, validated within the FACS cohort (Supplementary Figure 2E-F).

**Figure 2.**
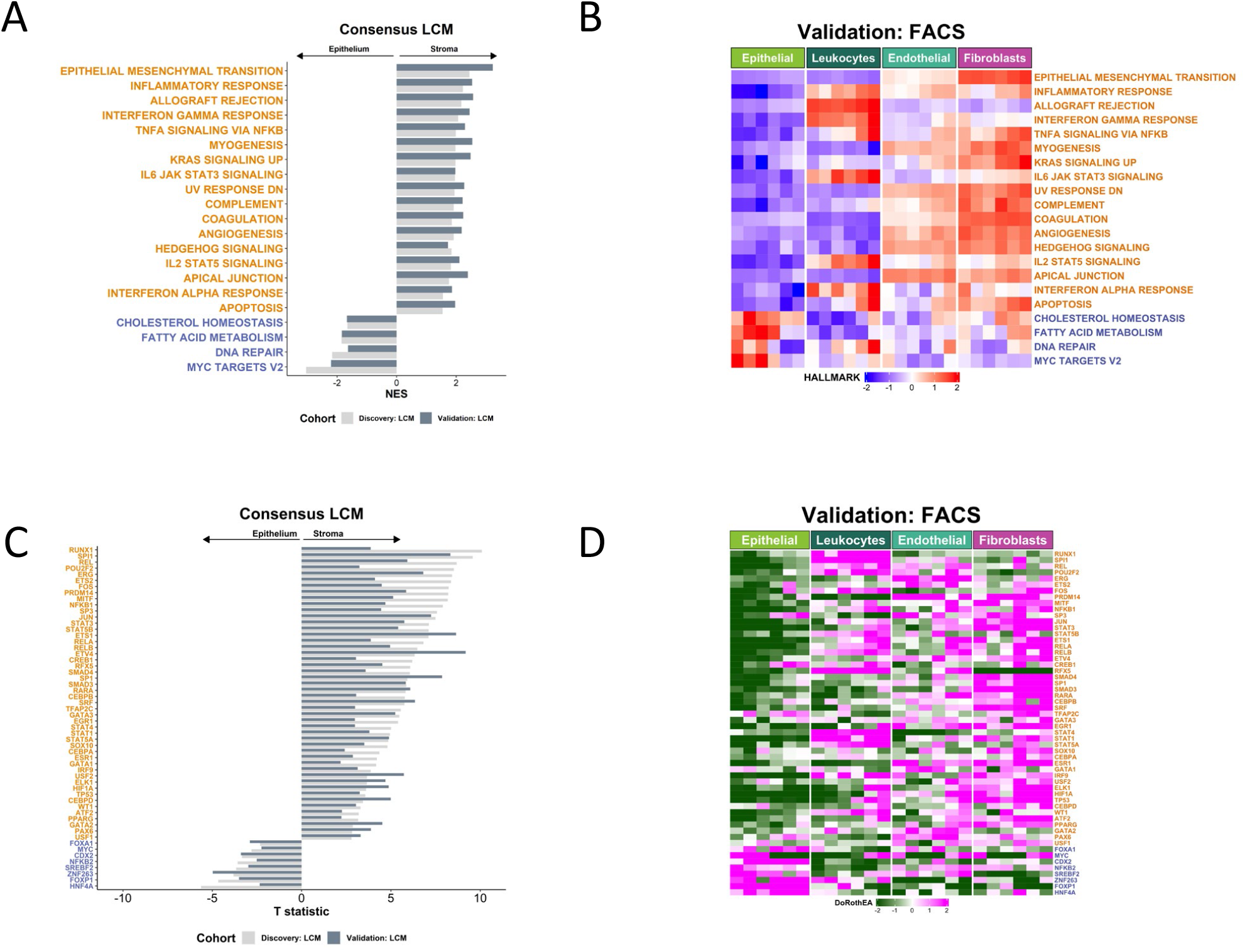
Stromal influence on widely used transcriptional signatures. **A** Gene set enrichment analysis (GSEA) of Hallmark gene sets in LCM discovery and validation cohorts. Only gene sets significantly and concordantly enriched in stroma or epithelium in both the discovery and validation cohorts are shown (adjusted p-value < 0.02). **B** Heatmap of single sample GSEA (ssGSEA) scores for the Hallmark gene sets in the FACS validation cohort samples. Only the gene sets significantly and concordantly enriched in stroma or epithelium in both the LCM discovery and validation cohorts are shown (adjusted p-value < 0.02). **C** Transcription factors whose activity was significantly and concordantly enriched in stroma or epithelium in both the LCM discovery and validation cohorts (p < 0.05). **D** Heatmap of the inferred activity scores for the same transcription factors in the FACS validation cohort. For all panels in Figure 2, gene sets/transcription factors with names/symbols coloured orange were significantly and consistently enriched in stroma in the LCM discovery and validation cohorts, whereas gene sets/transcription factors with names/symbols coloured blue were consistently and significantly enriched in epithelium in the LCM discovery and validation cohorts (gene sets: adjusted p-value < 0.02; transcription factors: p < 0.05).

We also observed similar confounding effects at the transcription factor (TF) activity level, when assessed using the n=118 defined regulons within the Dorothea algorithm.^22^ These analyses revealed the extent to which numerous seemingly epithelial-specific cancer-associated TFs are influenced by stromal content, across both LCM cohorts (Figure 2C); with n=48 TFs significantly activated in stromal components, compared to only n=8 TFs being significantly activated in the epithelium. As with the transcriptional signatures, when extended into the FACS purified populations, we observed a near identical overlap with the LCM findings and identified a number of lineage-specific associations (Figure 2D). Given the potential implications of the CRC findings described above, we next questioned if this was a pan-cancer phenomena, by performing the same analysis in LCM cohorts from breast cancer (BC), triple-negative breast cancer (TNBC), pancreatic ductal adenocarcinoma (PDAC), ovarian cancer (OvC) and prostate cancer (PrC). Despite some small organ-specific discrepancies in individual GESs, these analyses again revealed that the presence and extent of the confounding effect of the stroma is not CRC-specific, highlighting the potential for widespread biological misinterpretation of these signalling pathways across multiple cancer types (Supplementary Figure 2G-K).

### The ConfoundR resource enables estimation of stromal influence on transcriptional signatures simultaneously across multiple cancer types

Our findings of the presence of the stromal confounding effect across cancers, coupled with the widespread interest in biomarker/GES identification and application, motivated us to develop the online resource, ConfoundR (www.confoundr.qub.ac.uk). ConfoundR enables users, regardless of their bioinformatics skillset, to examine individual genes, combination of genes, and GES of interest to identify if they could be susceptible to the same stromal confounding issues we have identified in this study. This freely-available online resource enables users to interrogate the CRC, BC, TNBC, PDAC, OvC and PrC datasets through three analysis modules; gene expression boxplots, gene expression heatmaps and GSEA (Figure 3A).

**Figure 3.**
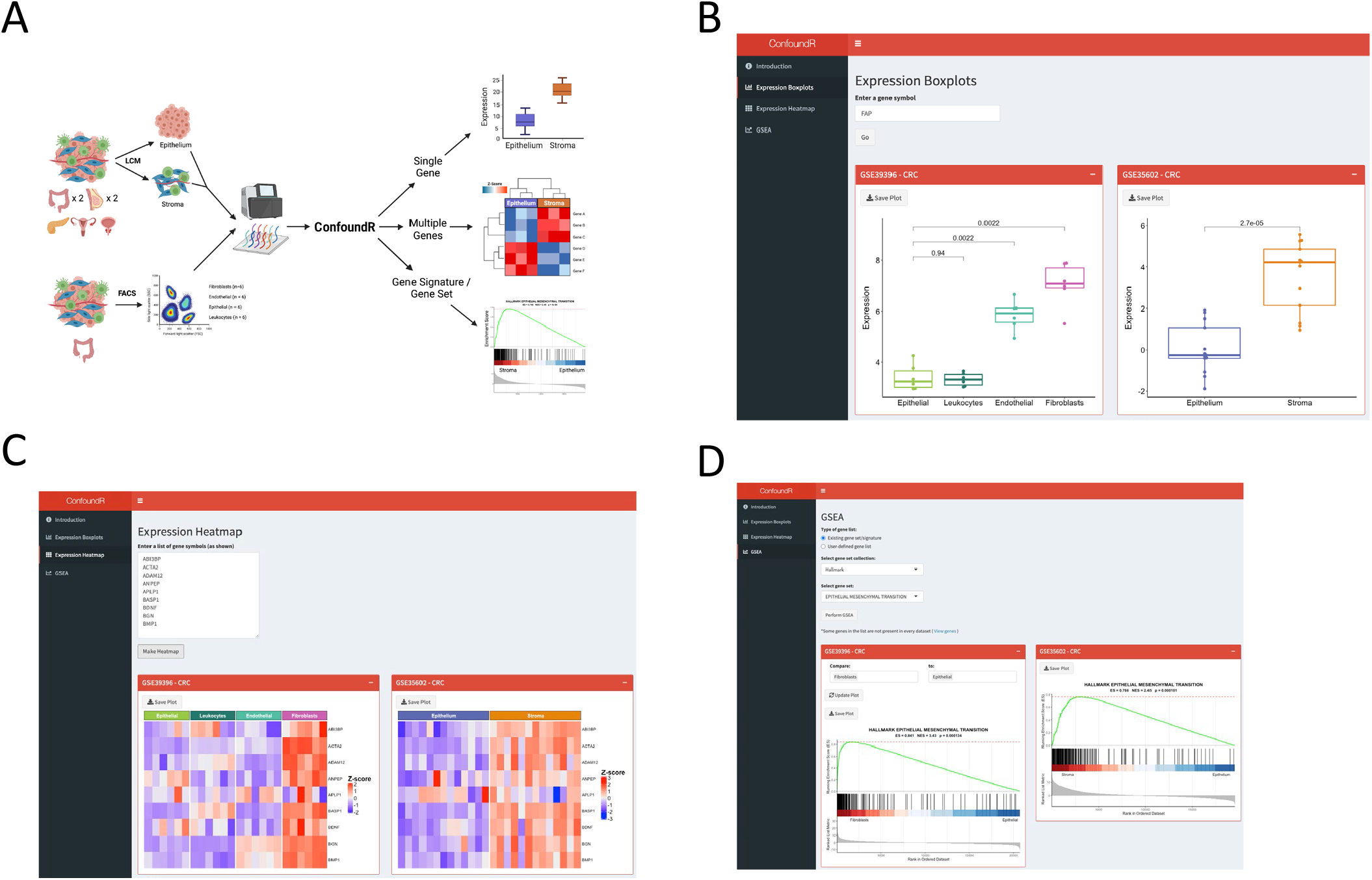
The ConfoundR resource enables stromal influence estimation in cancer tissue. **A** Schematic overview of the cohorts and analyses available within the ConfoundR app, accessible via https://confoundr.qub.ac.uk/. **B** Expression Boxplots analysis module of ConfoundR enabling the expression of a single gene to be compared between stroma and epithelium samples in each of the ConfoundR datasets. **C** Expression Heatmap analysis module of ConfoundR enabling the expression of multiple genes to be visually compared between stroma and epithelium samples in each of the ConfoundR datasets. **D** GSEA analysis module of ConfoundR allowing GSEA of existing gene sets from established gene set collections or custom user defined gene sets to be performed comparing stroma to epithelium in each of the ConfoundR datasets.

The boxplots module of ConfoundR allows gene expression comparisons of a single gene between epithelial samples and stroma samples in each dataset, by creating boxplots and providing accompanying p-values for Mann-Whitney U tests (Figure 3B). ConfoundR’s heatmap module allows expression levels of multiple genes to be visually compared between epithelial and stromal samples in each dataset using a heatmap (Figure 3C). Finally, the GSEA module of ConfoundR, enables the user to perform GSEA comparing stromal and epithelial samples in each dataset from established gene set collections; Hallmarks (n=50), KEGG (n=186), Reactome (n=1604), Biocarta (n=292), Pathway Interactions Database (PID; n=196). In addition, as many researchers will be interested in assessing their own bespoke or unpublished gene signatures, ConfoundR also provides the end-user with complete control to input and generate GSEA results from an unlimited number of custom gene sets (Figure 3D). To exemplify the utility of the ConfoundR resource, we examined the expression of the fibroblast activated protein (*FAP*) gene using the Expression Boxplots module, the expression of a subset of genes from the Hallmark EMT gene set using the Expression Heatmap module and the GSEA module to perform GSEA for the Hallmark EMT gene set (Figure 3B-D). The ConfoundR application provides all cancer researchers with a freely-available and novel resource to test the susceptibility of any gene, lists of genes or gene signatures to the stromal confounding phenomenon described in the present study.

### Application of findings to bulk CRC tumour data

To test these findings further in bulk tumour datasets, we utilised transcriptional data from n=356 primary tumours used in the FOCUS clinical trial (Figure 4A; GSE156915)^23^, alongside digital pathology-derived desmoplastic stromal percentage score derived from H&Es (desmoplastic stroma percentage [DS%]; detailed in Methods). We confirmed the previously-reported associations between CMS4 and stromal content are also observed in this tumour cohort (Supplementary Figure 3A) alongside strong correlation between our digital pathology assessments of stroma and the MCP fibroblast score (ρ=0.64, p<2.2e-16) (Supplementary Figure 3B) and ESTIMATE^24^ stromal score (ρ=0.73, p<2.2e-16) (Figure 4B). Using DS% to rank the tumour samples from low to high, we next assessed all the Hallmarks (Supplementary Figure 3C), alongside the subset of Hallmarks and TFs that were found to be significantly associated with stroma/epithelium in the LCM and FACS cohorts (Figure 4C and 4D), revealing a strikingly clear pattern that again indicates how widely the stromal components of a tumour can confound the interpretation of transcriptional signatures and TF activity scores in the bulk tumour setting.

**Figure 4.**
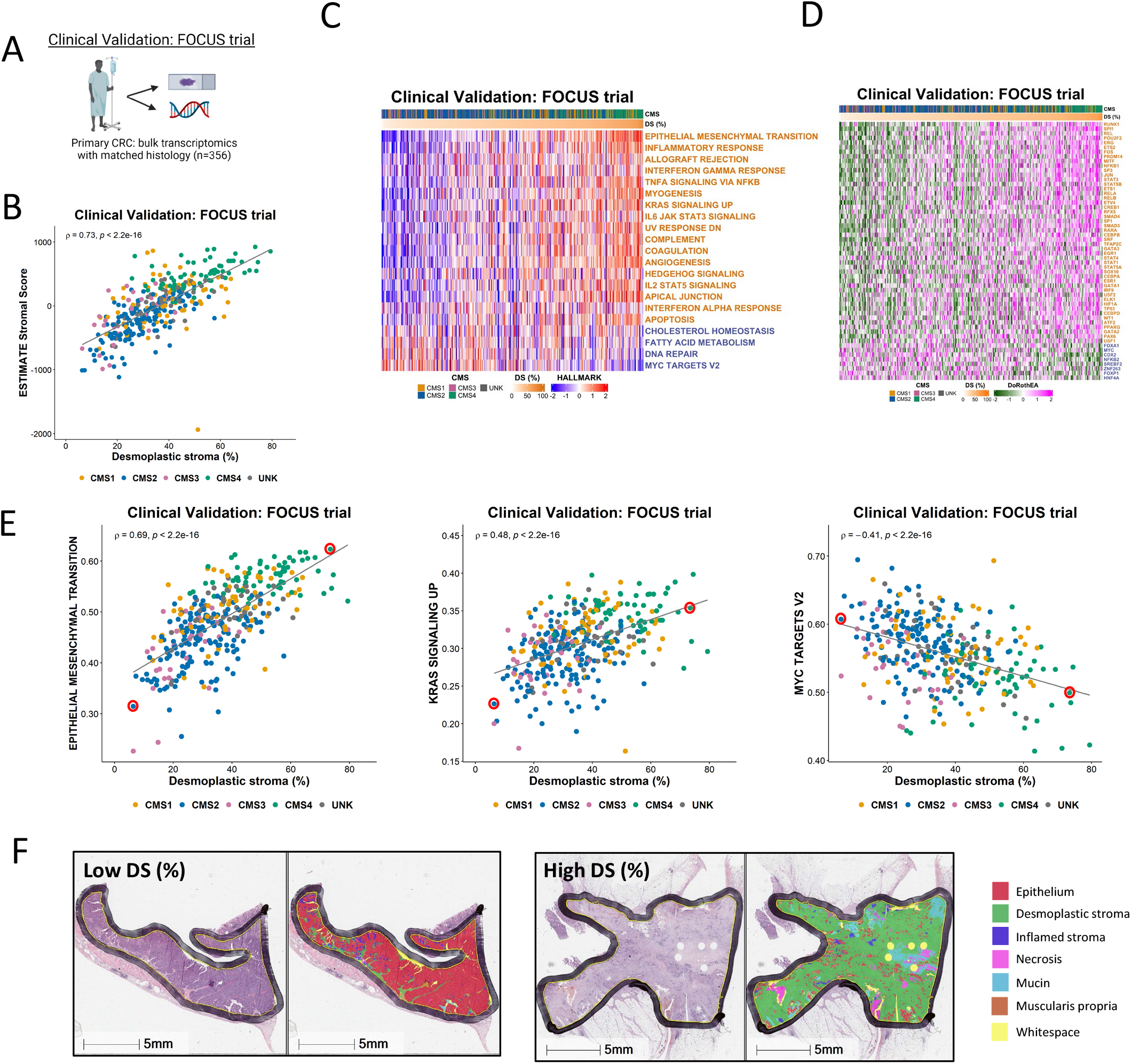
Application of findings to bulk CRC tumour data. **A** Schematic summary of the clinical validation dataset from the FOCUS clinical trial. **B** Scatterplot showing correlation between desmoplastic stroma percentage determined from H&E assessment and ESTIMATE Stromal Score determined by transcriptomic data in the FOCUS clinical trial samples (Spearman’s rho = 0.73, p < 2.2e-16), coloured by Consensus Molecular Subtype (CMS) calls (CMS1: n=62; CMS2: n=155; CMS3: n=29; CMS4: n=66; UNK: n=44). **C** Heatmap of ssGSEA scores for the Hallmark gene sets (identified in Figure 2 as significantly enriched in the stroma/epithelium in the LCM discovery and validation cohorts) for the FOCUS clinical trial samples. Samples ranked in order of desmoplastic stroma percentage (DS%) from lowest (left) to highest (right). Gene sets with names coloured orange were significantly enriched in stroma in the LCM discovery and LCM validation cohorts and gene sets with names coloured blue were significantly enriched in epithelium in the LCM discovery and LCM validation cohorts. **D** Heatmap of activity scores for transcription factors (identified as significantly enriched in the stroma/epithelium in the LCM discovery and validation cohorts) for the FOCUS clinical trial samples. Samples are arranged in order of desmoplastic stroma percentage (DS%) from lowest (left) to highest (right). Gene sets with names coloured orange were significantly enriched in stroma in the LCM discovery and LCM validation cohorts and gene sets with names coloured blue were significantly enriched in epithelium in the LCM discovery and LCM validation cohorts. **E** Scatterplots showing the correlation between desmoplastic stroma percentage determined from H&E and ssGSEA scores for the Epithelial Mesenchymal Transition (left) (Spearman’s rho = 0.69, p < 2.2e-16), KRAS Signalling Up (middle) (Spearman’s rho = 0.48, p < 2.2e-16) and MYC Targets V2 (right) (Spearman’s rho = -0.41, p < 2.2e-16) Hallmark gene sets. We identified two cases representative of low and high desmoplastic stromal percentage in each of these analyses (red circles). **F** H&E along with HALO mark-up for the representative low and high desmoplastic stromal percentage samples identified in E.

Throughout our analyses a number of individual signatures were consistently associated with the strongest confounding effects of stromal content, and therefore we selected these as specific exemplars related to DS%; namely the EPITHELIAL MESENCHYMAL TRANSITION (Spearman’s rho = 0.69, p < 2.2e-16), KRAS SIGNALLING UP (Spearman’s rho = 0.48, p < 2.2e-16) and MYC TARGETS V2 (Spearman’s rho = -0.41, p < 2.2e-16) (Figure 4E) Hallmark signatures. We identified two cases representative of low and high DS% in each of these analyses (Figure 4E, red circles) and assessed histological features according to H&Es with AI-guided tissue segmentation, which provided a visual confirmation that these Hallmark signatures are confounded by quantity of desmoplastic stroma across the tissue section (Figure 4F).

### Lineage-specific scRNAseq assessment of the Hallmarks EMT signature

Single cell RNA seq (scRNAseq) can be deployed to provide exceptional lineage-specific resolution in transcriptional studies, and this method has been used to great effect in the identification of tumour heterogeneity and phenotypic associations.^25^ To assess how far our findings extend in such data, we utilised a scRNAseq cohort derived from n=6 CRC primary tumours (Figure 5A; GSE144735), where across all regions at a single cell resolution the Hallmark EMT signature displays a significant enrichment in stromal cells compared to all other cell types (p <2.2e-16; Figure 5B) and in particular when comparing epithelial and stromal only (p <2.2e-16; Figure 5C). The highest EMT scoring epithelial cells only ever display an EMT gene expression signature score that reaches the lowest quartile of EMT signature score for stromal cells across all samples (Figure 5D). Based on these data, despite EMT signatures proving to be highly-tractable biomarkers of epithelial cells undergoing transitions when utilised in *in vitro*, pre-clinical or scRNAseq data, these data provide further proof that when applied to clinical samples, any EMT-related signature score, regardless of how well refined it is from pre-clinical models or scRNAseq data, becomes a definitive measurement of stromal content rather than epithelial to mesenchymal transition.

**Figure 5.**
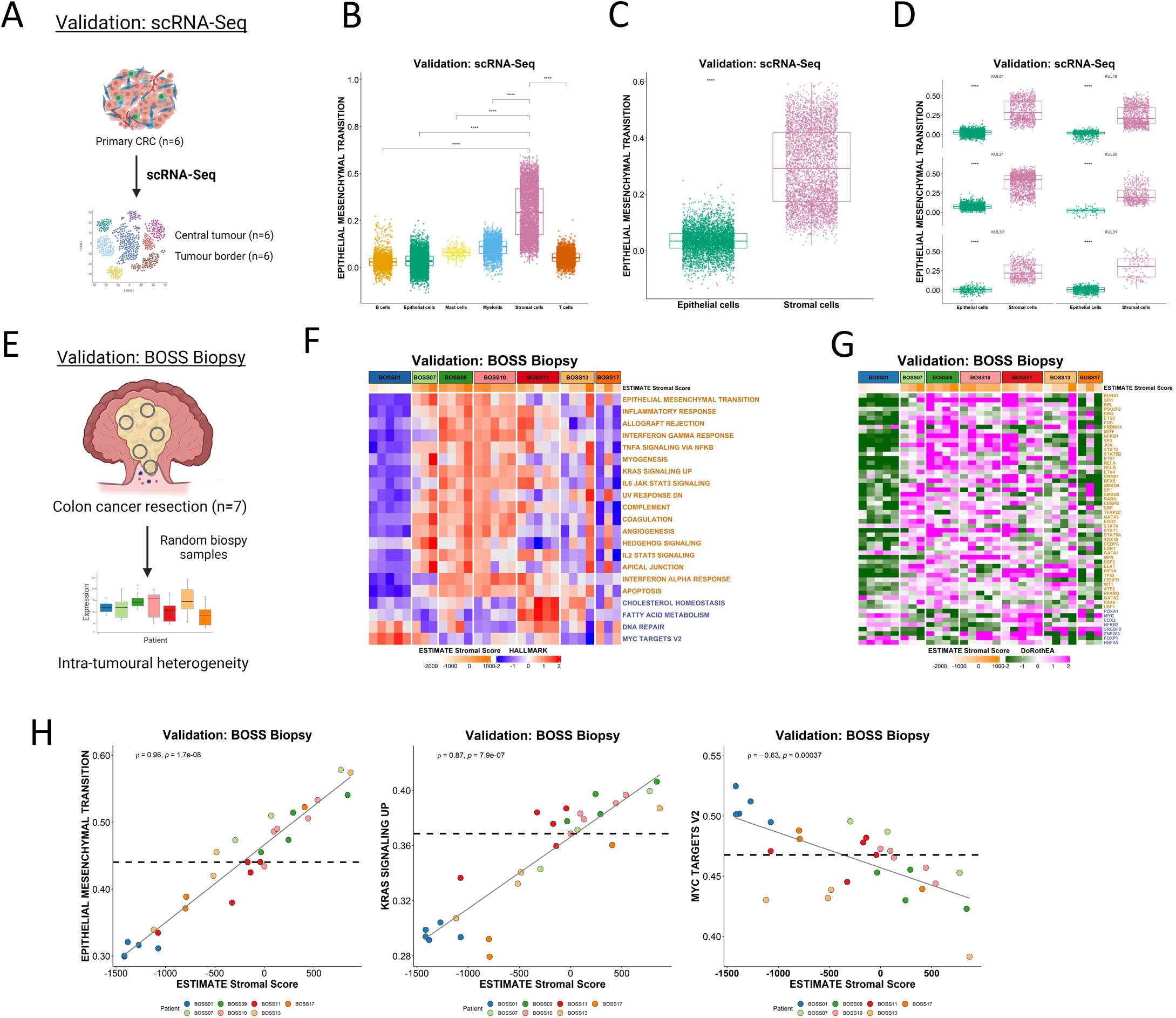
Single cell and multi-regional biopsy analyses. **A** Schematic of scRNA-Seq cohort derived from n=6 CRC primary tumours segregated into two regions for each tumour: the central tumour and tumour border. **B-D** Boxplots showing ssGSEA scores for the Hallmark Epithelial Mesenchymal Transition gene set **B** across the various cell types **C** and specifically between epithelial and stromal cells (from all six CRC tumours) in the scRNA-Seq dataset (p < 2.2×10-16; Wilcoxon test). **D** Comparison of ssGSEA scores for the Hallmark Epithelial Mesenchymal Transition gene set between epithelial and stromal cells in each primary CRC (n=6) in the scRNA-Seq dataset (all p < 2.2×10-16; Wilcoxon test). Epithelial cells are shown in green and stromal cells in pink. **E** Schematic overview of the BOSS Biopsy cohort consisting of colon cancer resections from seven patients each with up to n=5 multi-regional biopsy samples. **F-G** Heatmaps of **F** ssGSEA scores for the Hallmark gene sets and **G** transcription factor activity scores for the BOSS Biopsy samples. Samples are grouped according to patient of origin and the ESTIMATE StromalScore of each biopsy sample is indicated by the ESTIMATE StromalScore bar at the top of the heatmap. Only the gene sets/transcription factors significantly and concordantly enriched in stroma or epithelium in both the LCM discovery and LCM validation cohorts are shown (from Figure 2) (adjusted p-value < 0.02 – Hallmarks; p < 0.05 – transcription factors). Gene sets/transcription factors with names/symbols coloured orange were significantly enriched in stroma in the LCM discovery and LCM validation cohorts and gene sets/transcription factors with names/symbols coloured blue were significantly enriched in epithelium in the LCM discovery and LCM validation cohorts. **H** Scatterplots showing correlation between the ESTIMATE StromalScore and ssGSEA scores for the Hallmark Epithelial Mesenchymal Transition (left) (Spearman’s rho = 0.96, p = 1.7e-08), KRAS Signaling Up (middle) (Spearman’s rho = 0.87, p = 7.9e-07) and MYC Targets V2 (right) (Spearman’s rho = -0.63, p = 0.00037) gene sets. Samples are coloured by patient of origin.

### Multi-regional biopsy assessment

We next wished to test the potential clinical implications of these findings, in terms of patient misclassification, using the biopsy of surgical specimens (BOSS)^26^ cohort of n=7 primary colon tumour resections, where each patient tumour has bulk transcriptional profiles derived from up to n=5 multi-regional biopsies (Figure 5E). Application of ssGSEA for the Hallmarks revealed some signature- and patient-specific variations indicative of stromal-derived intratumoural heterogeneity. When assessed individually, all n=5 biopsy samples derived from patient BOSS01 display low expression of all n=17 Hallmarks and n=42 TFs we have previously associated with stroma, in line with this patient having a largely uniform epithelial-rich tumour (Figure 5F and 5G). However, for the remaining patient samples, particularly from patient BOSS11, BOSS13 and BOSS17, all displayed large variation in gene expression between their patient-matched biopsies for each of the stromal-associated n=17 Hallmark signatures and n=48 TFs (Figure 5F and 5G), suggesting that these tumours in particular displayed intratumoural heterogeneity in tumour stroma percentage. To test if the source of this intratumoural heterogeneity in Hallmark scores was due to variation in DS% content across biopsies, we assessed the individual ssGSEA signature scores correlated with the ESTIMATE stromal score which we previously confirmed as an accurate surrogate of DS% (Figure 4B). Remarkably, these analyses revealed the extent to which stromal content can accurately predict transcriptional signature scores regardless of the patient-of-origin. This was particularly evident for the signatures we have identified to be confounded by stromal content in our LCM, FACS and bulk tumour datasets, exemplified by positive correlation of EPITHELIAL MESENCHYMAL TRANSITION (Spearman’s rho = 0.96, p = 1.7e-08), KRAS SIGNALLING UP (Spearman’s rho = 0.87, p = 7.9e-07) signalling, alongside negative correlation with the MYC TARGETS V2 (Spearman’s rho = -0.63, p = 0.00037) signature (Figure 5H).

### Spatial transcriptomic confirms the confounding effects of the stroma

In this study, we have shown the potential for TSP to confound transcriptional signature scores, which in turn can result in misinterpretation of their meaning. Furthermore, analysis in the BOSS cohort also reveal the potential clinical implications of intratumoural stromal heterogeneity, which could result in patient misclassification, or indeed multiple conflicting classifications, when using GESs. To directly assess if spatial transcriptomics (ST) can alleviate some of the confounding variations in transcriptional signalling and inaccurate interpretation of findings when using bulk data, we profiled n=11 regions of a colon tumour sample using the GeoMx ST platform (Figure 6A). While the GeoMx Cancer Transcriptome Atlas gene panel employed was more limited (n=1825 core genes in total) when compared to the profiling in our other cohorts; we demonstrated that the reduced total number of genes still represent excellent surrogates for the whole transcriptome by assessing ssGSEA scores of the full signatures alongside the reduced genes available in the ST data. To test this further, using data from the FOCUS cohort, we observed excellent concordance in ssGSEA scores of the full MSigDB Hallmark EMT signature (ρ=0.95; n=200 genes) and MYC Targets V2 signature (ρ=0.75; n=58 genes), when assessed using the corresponding reduced signatures that were present on the GeoMx panel (n=81 genes and n=8 genes respectively) (Figure 6B, Supplementary Figure 4A). The ST platform provided the option to stratify our regions of interest into epithelium and stroma, using cytokeratin (pan-CK) staining, enabling calculation of total cellularity for each transcriptional pool. Using the ST data, we next performed ssGSEA using the Hallmarks we have previously shown to be most confounded by stroma, which again revealed the same general pattern across the n=17 stromal-associated and n=4 epithelial-associated signatures (Figure 6C). These findings were further confirmed when ST data from across the entire slide was pooled into two groups for pair-wise GSEA, PanCK- and PanCK+ (Supplementary Figure 4B), which again revealed a significant enrichment for the EMT Hallmark signalling cascade in the stromal (PanCK-) regions (Figure 6D). While bulk tumour datasets will remain an essential tool for statistical association studies, these data clearly highlight the need for the compartment and/or lineage-specific stratification, as afforded by ST, to ensure accurate biological interpretation of GESs.

**Figure 6.**
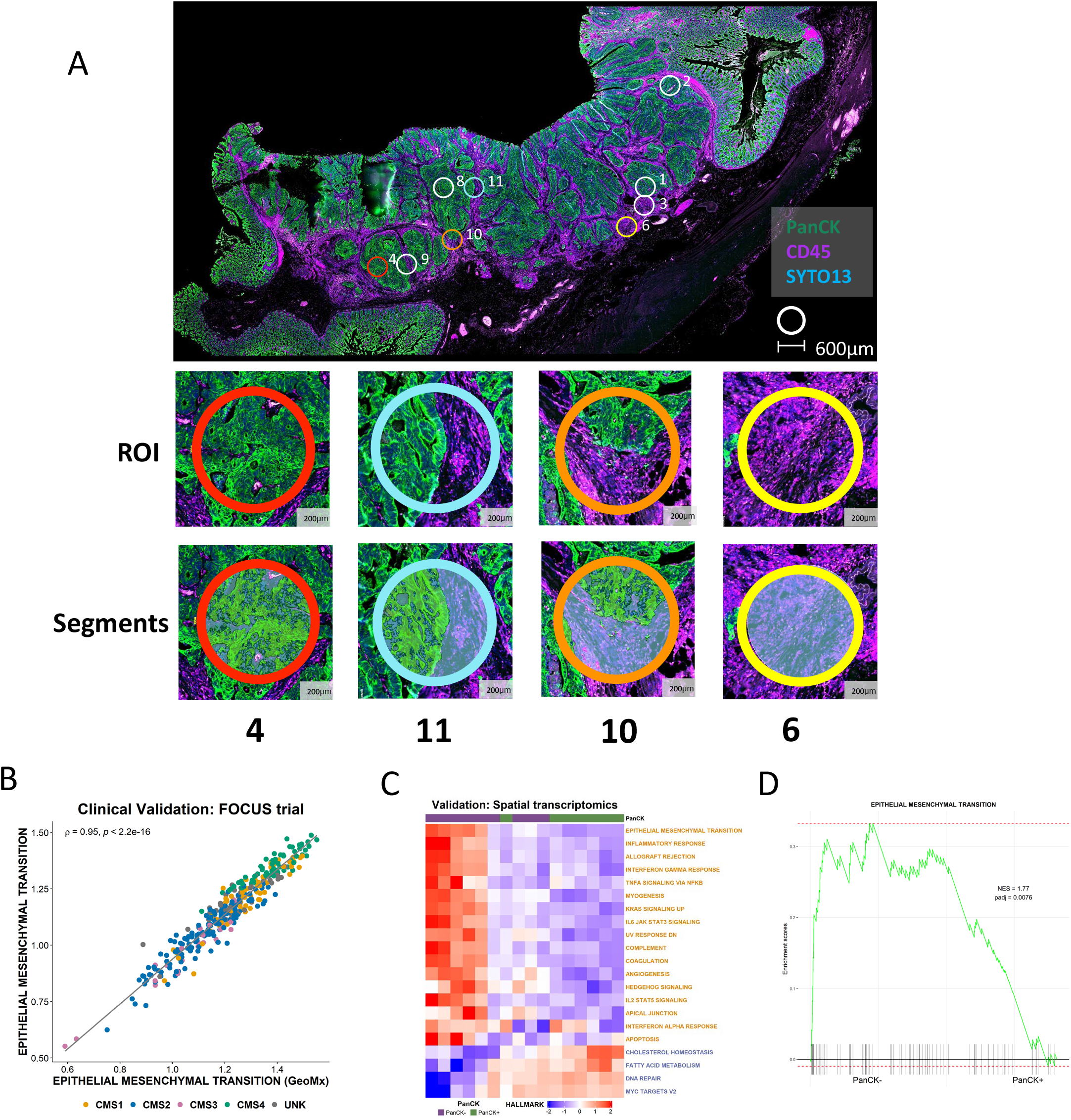
Spatial transcriptomic confirms the confounding effects of the stroma. **A** Whole slide image of colon cancer case selected for spatial transcriptomic analysis. The tissue was stained with Pan-Cytokeratin (PanCK) and CD45 with PanCK+ regions (green) identifying epithelium and CD45+ regions (purple) identifying immune components. Small circles indicate the regions of interest selected for spatial transcriptomic analysis; ROI 4: high epithelial content, ROI 11: mixed epithelial content, ROI 10 demonstrates a ROI with low epithelial content, ROI 6: no epithelial content. **B** Scatterplot showing the correlation between ssGSEA scores for the full Hallmark Epithelial Mesenchymal Transition gene set (n=200 genes) and the corresponding reduced GeoMx Epithelial Mesenchymal Transition gene set (n=81 genes) in the FOCUS clinical trial cohort (Spearman’s rho = 0.95). Samples coloured by Consensus Molecular Subtype (CMS) calls (CMS1: n=62; CMS2: n=155; CMS3: n=29; CMS4: n=66; UNK: n=44). **C** Heatmap of ssGSEA scores for the Hallmark gene sets for the PanCK+ (epithelium) (n=8) and PanCK-(stroma) (n=11) areas within the regions of interest. Only the Hallmark gene sets identified as significantly and concordantly enriched in stroma or epithelium in both the LCM discovery and LCM validation cohorts are shown (the GeoMx versions of these Hallmark gene sets were used). **D** GSEA comparing PanCK-areas (stroma) (n=11) to PanCK+ areas (epithelium) (n=8) for the Hallmark Epithelial Mesenchymal Transition gene set (GeoMx version).

## Discussion

In this study, we provide a comprehensive characterisation of the tumour transcriptome, stratified primarily into epithelium and stroma using LCM and spatial transcriptomics, alongside a more granular assessment of individual lineages using FACS and scRNAseq analysis. These analyses provide insight into the extent of the stroma’s contribution to some of the most widely-employed signatures in cancer research, and also the potential for biological misinterpretation of resulting data when extrapolating biology from preclinical models that do not contain a full tumour microenvironment. Furthermore, we can clearly show the potential clinical implications of these issues in terms of variable patient classification, both through the use of bulk tumour data (as demonstrated in this study from the FOCUS clinical trial) and through the use of multi-regional biopsies from primary resection material. To ensure that all users can benefit from the findings of this study, we have developed a user-friendly and freely-available resource, ConfoundR, which enables assessment of individual genes, pathways and bespoke signatures across a number of CRC, BC, TNBC, PDAC, OvC and PrC datasets.

Pre-clinical models, particularly epithelial-rich systems where near-complete control over lineage purity and environmental conditions can be achieved and reproduced, represent ideal systems to develop transcriptional signatures that correlate with phenotypes of interest. We and others have previously highlighted discrepancies between the nomenclature used for such published signatures,^4,27^ when named to reflect the phenotypes they characterise *in vitro*, and the actual biology they can represent when applied to bulk tumour datasets. This is primarily due to the fact that while lineage purity is fixed in such pre-clinical models, a tumour mass is composed of a milieu of lineages,^13^ the proportions of which are most often unknown at the time of processing for bulk transcriptomic profiling. This becomes particularly problematic for signatures and biomarkers that are developed to characterise mesenchymal traits, as although they will track with precise epithelial biology in *in vitro* systems, when applied to tumour data they are more likely to become highly-accurate tumour stromal percentage estimates, rather than measures of subtle epithelial transitions.^5,6^ Our current study addresses the importance of also ensuring that the transcriptional signatures faithfully represent the same biology during forward and reverse translation studies, and are not undermined by changes in conditions between clinical and preclinical settings. As such, the most faithful alignment of biological traits between models and clinical samples should be based on deeper phenotyping assessments, that incorporate histology and lineage-specific assessments in addition to transcriptional signatures.

The contribution of the stroma to the cancer transcriptome has been the subject of numerous studies, highlighted by Isella and colleagues^18,28^ who identified a set of 4,434 genes present in cancer datasets but exclusively expressed by the tumour stroma. While the nomenclature used in each of the signatures tested may lead researchers to conclude that the biology of that pathway is elevated, data presented here clearly demonstrates that when these otherwise mechanistic-driven signatures are composed of genes expressed at higher levels in stromal cells, the signatures themselves become surrogate markers, to different extents, of the tumour stroma percentage within a bulk tumour sample. Given the extent to which the stroma appears to confound biological interpretation of the thousands of signatures and TFs we have assessed in this study, there may be a significant body of research published that has inadvertently derived conclusions based on inaccurate interpretations of results.

Data presented here do not challenge the use or value of using signatures to interpret data from bulk tumours, but present unambiguous intelligence around the caution that should be applied when ***interpreting*** what these signature scores can reflect, despite what the signature name suggests. An inaccurate biological conclusion in itself will not have major consequences for the statistical correlation associated with a signature, however it is highly likely that inaccurate biological interpretations of such results are being used as the basis for ongoing clinical therapeutic developments, which themselves are also potentially being tested in models that bear little relevance to the patient tumour samples they are derived from. This is particularly important, as most users are reliant on utilising existing molecular datasets, where there is no control over the initial profiling steps, or indeed representative histological images that precisely align to the tumour region used for nucleic acid extraction. As such, while the signatures presented here represent large collections of experimentally-validated genes associated with specific phenotypes or biological cascades, our findings support the conclusion that widespread misconceptions exist when interpreting the meaning underlying transcriptional signatures in tumour studies, given the discordance between their development and application.

Genes can have many functions, and in some cases the genes that strongly demark a specific phenotype in a pre-clinical model system can also be expressed and perform entirely different mechanistic signalling roles in stromal lineages that make up a TME. If the magnitude of expression of these genes is low in the stromal/immune lineages, this may cause minimal impact when interpreting the precise nature of the transcriptional signature in bulk tumour data. However, if the genes within these signatures are expressed at higher levels in non-epithelial lineages, they can become strong surrogate markers for levels of TME components, rather than reflecting any of the mechanistic biology that they were designed to identify. It is this potential for misinterpretation that our paper wishes to highlight, thereby enabling researchers to assess the potential for their mechanistic signature of choice to be confounded in bulk tumour data using our ConfoundR resource. This will hopefully allow researchers to make a more informed interpretation of the true biology underlying the signature in these tumour samples, as compared to the controlled pre-clinical system it was developed in.

While the use of scRNAseq analysis can provide high quality lineage-specific transcriptional data, this comes at the expense of spatial information.^29^ Conversely, spatial transcriptomics can regionalise transcriptional signalling but lose the single cell resolution.^30^ While both approaches, individually or in combination,^31,32^ have revolutionised the field of transcriptional profiling, the use of bulk transcriptomics datasets available in publicly-accessible databases like TCGA and GEO, remain the mainstay for alignment of transcriptional signature to clinical outcome data for prognostic assessment and mechanistic/biological interpretation. It is likely that with reducing costs and expansion of technologies, the generation of tumour-matched scRNAseq, spatial and bulk cohorts in both clinical samples and pre-clinical models will become more routine in future, and at some point may supersede that of existing bulk data. However, this is unlikely to be in the immediate future and as such the findings of this study, and the unique ConfoundR application we have made publicly-available, will represent an important resource to ensure that translational researchers can more accurately interpret the information underpinning the transcriptional biomarker(s) used to stratify patient samples and inform cancer care.

## Methods

### Datasets

When publicly available, the data were assessed via GEO and the processed data matrix downloaded. All array data were collapsed using the collapseRows function within WGCNA (v1.70-3) R package. In the case of duplicated genes, the probe with the highest mean expression across all samples was used and those genes with no expression across the dataset were removed. All GEO datasets are available via https://www.ncbi.nlm.nih.gov/geo/ using GSE codes below:

*Discovery: LCM* GSE35602; Matched epithelium and stroma from thirteen colorectal tumours transcriptionally profiled using Aligent array. *Validation: LCM* GSE31279; Matched epithelium and stroma from eight colorectal tumours (with both compartments) transcriptionally profiled using Illumina sentrix-8 chip. *Validation: FACS* GSE39396; Six CRC tumours were FACS into four cell populations: epithelial cells (EPCAM+), leukocytes (CD45+), fibroblasts (FAP+) and endothelial cells (CD31+). *Validation: Breast Cancer* GSE14548; Matched LCM epithelium and stroma from nine invasive ductal carcinomas transcriptionally profiled using the Affymetrix Human X3P Array. *Validation: TNBC* GSE81838; Matched LCM epithelium and stroma from ten triple negative breast cancers transcriptionally profiled using the Affymetrix Human Gene 1.0 ST Array. *Validation: PDAC* GSE164665; Matched LCM epithelium and stroma from nineteen PDACs transcriptionally profiled by Illumina NextSeq 500. *Validation: Ovarian Cancer* GSE9899; Matched LCM epithelium and stroma from five ovarian tumours transcriptionally profiled using the Affymetrix Human Genome U133 Plus 2.0 Array. *Validation: Prostate Cancer* GSE97284; Matched LCM epithelium and stroma from 25 prostate tumours, of which 12 were low grade (Gleason 3+3) and 13 were high grade (Gleason 8 or above) transcriptionally profiled using the Affymetrix Human Gene 1.0 ST Array. *Clinical Validation: FOCUS trial* GSE156915; The UK Medical Research Council FOCUS [<u>F</u>luorouracil, <u>O</u>xaliplatin and <u>C</u>PT11 (irinotecan)] trial involving stage IV patients with primary CRC resection transcriptionally profiled on Almac Xcell chip, only those with matched histology remained for analysis (n=356). *Validation: scRNAseq* GSE144735; The border and central tumour from six colorectal patients were single cell sequenced. Count matrix was aligned to annotation file within Partek Genomics Suite. Genes with an expression <1 in at least 99.9% of cells were removed. Data was normalised by counts per million, +1 and log2 transformed. *Validation: BOSS biopsy* GSE85043; Multiple biopsies obtained with different regions of seven CRC resection specimens. Profiled on Affymetrix array.

### GeoMx

#### Nanostring GeoMx Digital Spatial Profiling

To further characterise differences in transcriptomic expression between TME and tumour epithelium, a formalin-fixed paraffin-embedded (FFPE) section of archival resected colon cancer was selected for analysis on the Nanostring GeoMx Digital Spatial Profiler (DSP). This platform enables the characterisation of user-selected topographic Regions of Interest (ROI) from immunofluorescently (IF) stained FFPE tissue.The GeoMx instrument achieves RNA profiling *in situ* hybridization by employing DNA oligonucleotide probes designed to bind mRNA targets. From 5′ to 3′, they comprise a 35-to 50-nucleotide target complementary sequence, an ultraviolet (UV) photocleavable linker and a 66-nucleotide indexing oligonucleotide sequence containing a unique molecular identifier (UMI), RNA ID sequence and primer binding sites. Up to 10 RNA detection probes were designed per target mRNA. In summary, the instrument employs UV light to cleave the UV-sensitive probes leading to release of the hybridised barcodes.

#### Slide preparation including hybridisation of tissue with UV-photocleavable probes

The DSP procedure has previously been described in detail by Merritt et al (1). The 5-µm FFPE tissue section was mounted on positively charged Superfrost glass slide (Thermo Fisher Scientific) and baked for 30 mins at 60 °C. The tissue was dewaxed, hydrated and treated with 1μg/ml Proteinase K (Thermo Fisher Scientific, AM2546) for 15 minutes before undergoing heat-induced epitope retrieval (HIER) on a Leica BOND Autostainer (pH 9.0, ER2 at 100°C) for 20 minutes. The slide was immediately stored in 1X PBS (PBS: Invitrogen, AM9625). Hybridisation with a pre-designed Cancer Transcriptome Atlas (CTA) panel of antibodies corresponding to 1,825 genes (Nanostring) was performed according to the manufacturer’s protocol (2). 100μL of the RNA probe mix (CTA panel) was mixed with 800μL of Buffer R (Nanostring) and 100μL of DEPC-treated water. Each tissue was covered with 200μL of hybridisation solution and a HybriSlip(tm) cover (Thermo Fisher Scientific) before overnight incubation in a hybridisation oven at 37 °C for at least 16 hours. The slide was then dipped in 2X SSC-T and washed twice with a 1:1 ratio of 100% deionized formamide (Ambion) and 4X SSC (Sigma) at 37°C for 25 minutes each.

The GeoMx DSP is capable of capturing four channels (FITC/525nm, Cy3/568nm, Texas Red/615nm and Cy5/666nm) for the detection of up to four customisable IF morphology markers for each tissue (1). One channel (FITC/525nm) is reserved for the nuclear stain (SYTO13). The slides were blocked with Buffer W (Nanostring) for 30 minutes at RT before incubation with TME RNA Morphology Marker kit (Nanostring) for 1 hour at RT. This consisted of fluorescently conjugated Syto13, Pan-Cytokeratin (PanCK) and CD45 antibodies were used to stain the tissue to identify nuclei, tumour epithelium and the immune components respectively. Slides were then stored at 4°C in SSC before being loaded on the GeoMx DSP instrument for ROI selection and collection.

#### Region selection and collection

The whole slide was imaged at 20x magnification using the GeoMx DSP with the integrated software suite then used to select 300-600um diameter ROIs from which the instrument focuses UV light (385nm), to cleave the UV-sensitive probes with the subsequent release of the hybridised barcodes. 11 ROIs corresponding to epithelial tumour centre, abundant TME regions and regions representing an interface between tumour and TME and were selected. The DSP software enabled Areas of Interest (AOI) contained in individual ROIs to be defined and selected. Firstly segments containing PanCK+ IF signal were masked for tumour epithelium and extracted, then the complementary inverse segments (PanCK-) was masked and captured corresponding to the TME. Once AOIs were defined, then exposed UV light, the indexing oligonucleotides, were collected with a microcapillary and deposited in a 96-well plate prior to sequencing. The oligonucleotides were dried overnight and subsequently resuspended using 10μl of DEPC-treated water

#### Library Preparation and NGS Sequencing

Sequencing libraries were generated by PCR from the photo-released indexing oligos and AOI-specific Illumina adapter sequences, and unique i5 and i7 sample indices were added. Each PCR reaction used 4µl of each collection sample added to the corresponding well of a new 96-well PCR plate containing the GeoMx Seq Code primers (Nanostring) and 1X PCR Master Mix (Nanostring). The PCR plate was incubated in a thermocycler with the programme specified by the manufacturer. The PCR products were then centrifuged and pooled (4μl each) into one 1.5 mL eppendorf to create a library. The library was purified twice using AMPure XP system (Beckman Coulter). The purified library was resuspended in Elution Buffer (10mM Tris-HCl with 0.05% Tween-20, pH 8.0) before undergoing quality check using an Agilent Bioanalyser.

The purified library underwent Next Generation Sequencing (NGS) using an Illumina NextSeq 550 (Glasgow Polyomics). Recommended sequencing parameters were followed in generation of FASTQ files, dual-indexing, paired-end reads and including a 5% PhiX spike-in. Sequencing depth was determined by total ROI area (µm^2^) multiplied by sequencing depth factor (30 for CTA panel) as per manufacturer’s instructions. H&Es from Focus (N=356) were scanned at high resolution on an Aperio scanner at a total magnification of 20X. Tissue segmentation was run on H&E images by deep convoluted neural net using the HALO platform (Indica Labs). Supervised training had been performed using >1,500 tissue areas from four CRC cohorts. Counts of single cells were utilised to assess the proportion of desmoplastic stroma compared to total cell counts.

#### Digital histology scoring

H&Es from Focus (N=356) were scanned at high resolution on an Aperio scanner at a total magnification of 20X. Tissue segmentation was run on H&E images by deep convoluted neural net using the HALO platform (Indica Labs). Supervised training had been performed using >1,500 tissue areas from four CRC cohorts. Counts of single cells were utilised to assess the proportion of desmoplastic stroma compared to total cell counts.

#### Data analysis

The FASTQ files generated were converted into Digital Count Conversion (DCC) files using the GeoMx NGS pipeline on the Illumina BaseSpace platform. The DCC files were uploaded onto the GeoMx DSP analysis suite (Nanostring), where they underwent quality control, filtering, Q3 normalisation and background correction. Data were then downloaded from the GeoMx instrument and loaded in to RStudio (v1.2.1335) using R build version 4.1.1.

### MCP

MCPcounter (v1.2.0) R package was used to generate scores for 10 cell populations. ***CMS***. Consensus Molecular Subtype classification utilised ‘classifyCMS.SSP’ function within the CMSclassifier (v1.0.0) R package and the CMScaller (v2.0.1) R package. ***ssGSEA***. Single sample gene set enrichment analysis was performed using gsva (v1.38.2) R package with the following non-default settings: min.sz=5, verbose = TRUE, method = ‘ssgsea’, on the HALLMARK, Gene Ontology: Biological Processes and the KEGG genesets. ESCAPE (v1.4) R package was utilised to generated single sample scores for the HALLMARK pathways within the single cell cohort using the ‘enrichIt’ function: groups= 1000, cores=2. ***GSEA***. Pair-wise gene set enrichment analysis was performed using fgsea (v1.16.0) R package (minSize=1, maxSize=Inf, nperm=10000), on the HALLMARK, Gene Ontology: Biological Processes and the KEGG genesets accessed via msigdb (v7.4.1). Within the FOCUS validation dataset a median split of DS was used for comparison, followed by DGEA. ***DoRothEA***. Transcriptional factor activity was assessed using dorothea (v1.2.2) R package, within the ‘run_viper’ function (filtered for high confidence regulons). Within LCM cohorts, P <0.05 was considered significant to obtain consensus LCM TFs. Plots in subsequent cohorts include all TF, regardless of significance. **ESTIMATE**. estimate (v1.0.13) R package was used to generate stromal and immune scores. R studio (v1.3.1073), R (v4.0) used for all analysis. All heatmaps were plotted using ComplexHeatmap and all additional plots using ggplot2.

### ConfoundR Shiny application

#### Development and Access

The ConfoundR application was created using R version 4.1.2 in combination with the R package shiny (RRID:SCR_001626; version 1.7.1) and is running on the Shiny Server (version 1.5.17) hosted on the Queen’s University Belfast virtual server (CentOS 7, 64-bit, Intel Xeon Gold E5-2660 v3 @2.60 GHZ, 16 Core). ConfoundR is accessible at https://confoundr.qub.ac.uk. **Datasets**. The datasets used in the ConfoundR application are described above, along with the pre-processing methods applied to each dataset.

#### Expression Boxplots

The Expression Boxplots module allows the user to enter the gene symbol for a single gene into the input box. Boxplots in the Expression Boxplots module are created using ggplot2 (RRID:SCR_014601; version 3.3.5) and Mann-Whitney U tests are performed using the stat_compare_means function from the ggpubr package (RRID:SCR_021139; version 0.4.0) with method = “wilcox.test”. Boxplots for the PDAC dataset (GSE164665) are plotted using normalised counts calculated by DESeq2 (RRID:SCR_015687; version 1.34.0), using the size factors calculated by the estimateSizeFactors function, accessed via the counts function with normalized = TRUE. Plots for each of the datasets can be downloaded in png format using the Save Plot button.

#### Expression Heatmap

The Expression Heatmap module enables users to enter a list of gene symbols with each gene symbol on a new line. For the RNA-Seq dataset, variance stabilising transformed counts, calculated using the vst function (blind = FALSE), from the DESeq2 package (RRID:SCR_015687; version 1.34.0), are used as the gene expression values for samples. The gene expression values for each user selected gene in each dataset are converted to Z-scores using the scale function (center = TRUE, scale = TRUE) prior to plotting heatmaps. Heatmaps of the gene expression Z-scores are plotted using the ComplexHeatmap package (RRID:SCR_017270; version 2.10.0) with the samples grouped by the respective cell/tissue types to aid visual comparison between groups.

### GSEA

The GSEA module enables users to select an existing gene set from established gene set collections using dropdown menus or to enter a custom user-defined gene set by entering a list of gene symbols with each symbol on a new line. The existing gene sets available to the user are the Hallmark, KEGG, Reactome, Biocarta and Pathway Interactions Database (PID) gene sets as curated by the Molecular Signatures Database (RRID:SCR_016863) and accessed via the msigdbr package (version 7.4.1).

In order to perform pre-ranked GSEA, differential analysis is performed for each of the datasets, comparing stromal samples to epithelial samples. For the GSE39396 dataset the user can specify the cell types (epithelial, leukocytes, endothelial, fibroblasts) to compare using the input boxes provided. Differential analysis is performed using limma (RRID:SCR_010943; version 3.50.0) for microarray datasets (GSE39396, GSE35602, GSE31279, GSE81838, GSE14548, GSE9899, GSE97284) and using DESeq2 (RRID:SCR_015687; version 1.34.0) for the RNA-seq dataset (GSE164665). Following differential analysis, genes are ranked according to the t-statistic (limma) or Wald statistic (DESeq2). Pre-ranked GSEA is performed by the GSEA function from the clusterProfiler package (RRID:SCR_016884, version 4.2.1) using the fgseaSimple method with 10,000 permutations (by = “fgsea”, nPerm = 10000) and a random seed of 123. Plots of GSEA results are produced using a modified version of the gseaplot2 function from the enrichplot package.

### Packages used

The ConfoundR app uses the following R packages: shiny (v1.7.1), shinydashboard (v0.7.2), dashboardthemes (v1.1.5), shinyFeedback (v0.4.0), shinybusy (v0.2.2), shinyccssloaders (v1.0.0), msigdbr (v7.4.1), ggplot2 (v3.3.5), cowplot(v1.1.1), ggbeeswarm (v0.6.0), ggpubr (v0.4.0), ComplexHeatmap (v2.10.0), limma (v3.50.0), DESeq2 (v1.34.0), clusterProfiler (v4.2.1), fgsea (v1.20.0), enrichplot (v1.14.1) and RColorBrewer (v1.1-2).

### Code availability

The source code for the ConfoundR app is available at https://www.github.com/Dunne-Group/ConfoundR. All scripts to perform the analyses outlined in this paper are available on our lab website www.Dunne-Lab.com.

