## Supplementary Figures for "Biological misinterpretation of transcriptional signatures in tumour samples can unknowingly undermine mechanistic understanding and faithful alignment with preclinical data"

#### Supplementary Figure 1

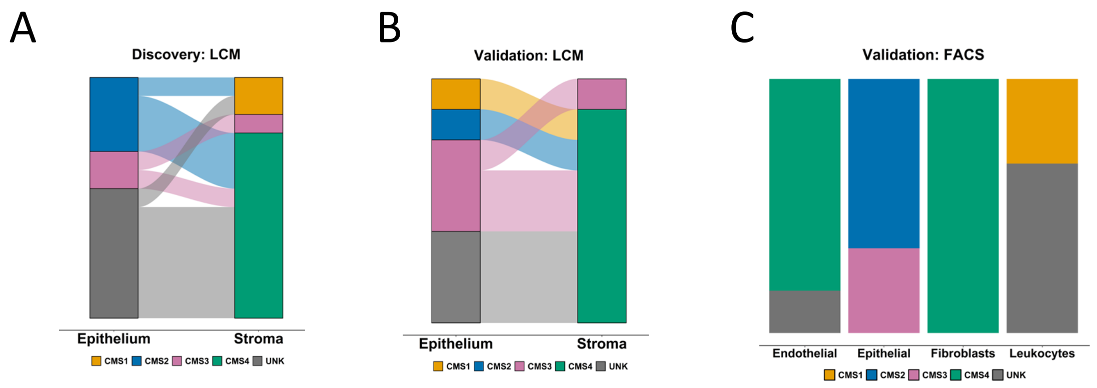


**Supplementary Figure 1**

**A** CMS calls (using CMScaller) for the matched epithelium and stroma samples in the LCM discovery cohort. **B** CMS calls (using CMScaller) for the matched epithelium and stroma samples in the LCM validation cohort. **C** CMS calls (using CMScaller) for the four lineages in the FACS validation cohort.

### Supplementary Figure 2

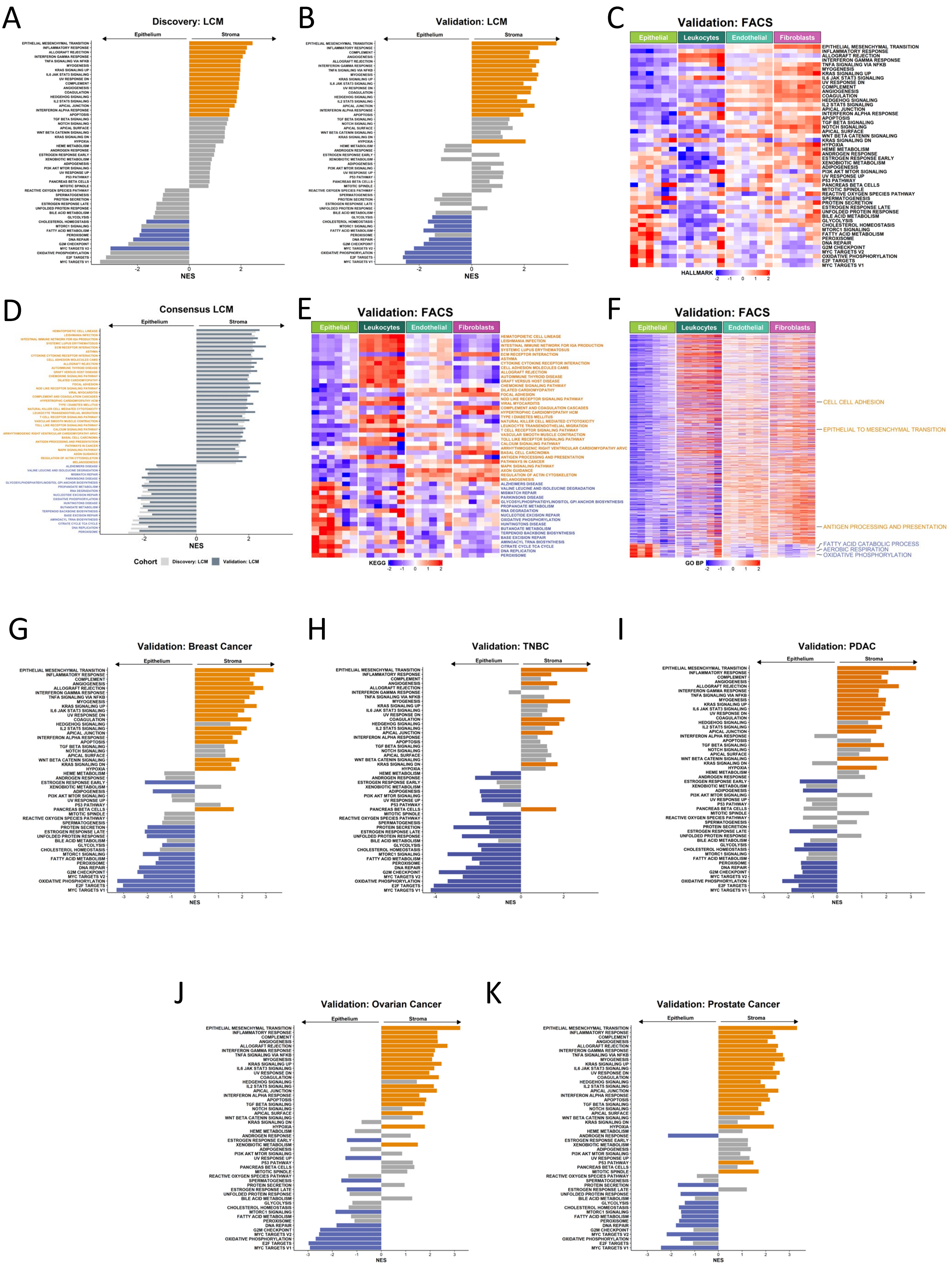

**Supplementary Figure 2**

**A** GSEA of Hallmark gene sets comparing stroma to epithelium samples in the LCM discovery cohort. Orange bars indicate gene sets significantly enriched in stroma, while blue bars indicate gene sets significantly enriched in epithelium (adjusted p-value < 0.02). **B** GSEA of Hallmark gene sets comparing stroma to epithelium samples in the LCM validation cohort. Orange bars indicate gene sets significantly enriched in stroma, while blue bars indicate gene sets significantly enriched in epithelium (adjusted p-value < 0.02). Gene sets are arranged in the same order as in the LCM discovery cohort (Supplementary Figure 2A). **C** Heatmap of ssGSEA scores for the Hallmark gene sets in the FACS validation cohort samples. Gene sets are arranged in same order as in the LCM discovery cohort GSEA (Supplementary Figure 2A). **D** GSEA of KEGG pathway gene sets comparing stroma to epithelium samples in the LCM discovery and validation cohorts. Gene sets with names coloured orange were significantly enriched in stroma in the LCM discovery and LCM validation cohorts and gene sets with names coloured blue were significantly enriched in epithelium in the LCM discovery and LCM validation cohorts. Only the gene sets significantly and concordantly enriched in stroma or epithelium in both the LCM discovery and LCM validation cohorts are shown (adjusted p-value < 0.02). **E** Heatmap of ssGSEA scores for the KEGG pathway gene sets in the FACS validation cohort samples. Only the gene sets significantly and concordantly enriched in stroma or epithelium in both the LCM discovery and LCM validation cohorts are shown (adjusted p-value < 0.02). Gene sets with names coloured orange were significantly enriched in stroma in the LCM discovery and LCM validation cohorts and gene sets with names coloured blue were significantly enriched in epithelium in the LCM discovery and LCM validation cohorts. Gene sets are arranged in the same order as in the LCM discovery cohort GSEA result (Supplementary Figure 2D). **F** Heatmap of ssGSEA scores for the Gene Ontology Biological Processes (GO BP) gene sets in the FACS validation cohort samples. Only the gene sets significantly and concordantly enriched in stroma or epithelium in both the LCM discovery and LCM validation cohorts are shown (adjusted p-value < 0.02) (Supplementary Table 1). **G-K** GSEA of Hallmark gene sets comparing stroma to epithelium samples in the **G** breast cancer, **H** triple negative breast cancer (TNBC), **I** pancreatic ductal adenocarcinoma (PDAC), **J** ovarian cancer and **K** prostate cancer cohorts. Gene sets are arranged in in the same order as in the LCM discovery cohort GSEA result (Supplementary Figure 2A).

#### Supplementary Figure 3

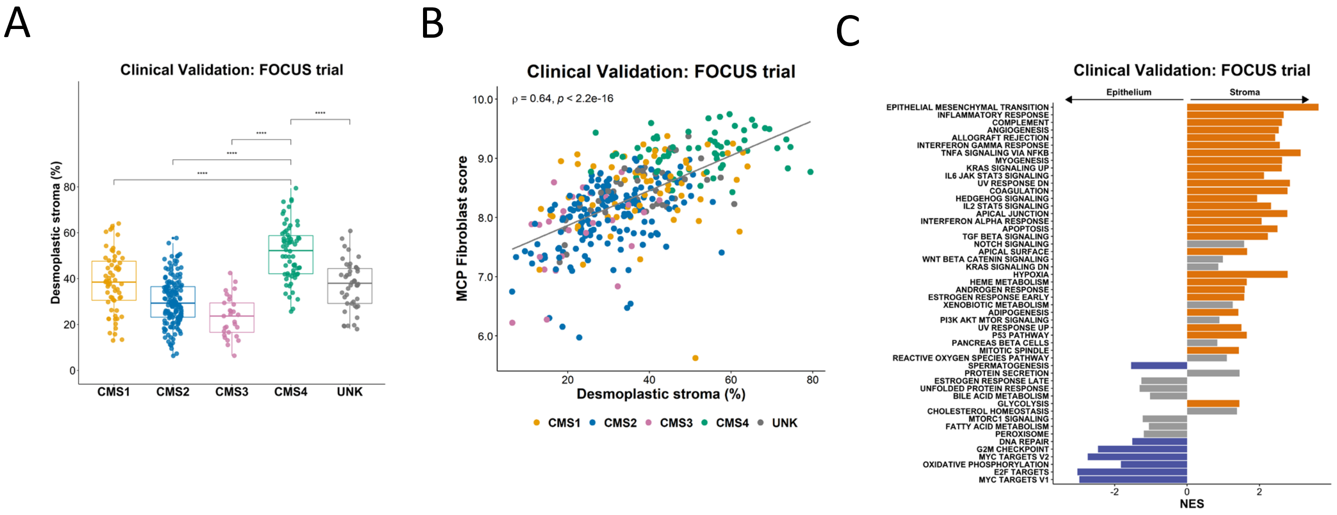


#### Supplementary Figure 3

**A** Boxplots showing desmoplastic stroma percentage across Consensus Molecular Subtype (CMS) calls (CMS1: n=62; CMS2: n=155; CMS3: n=29; CMS4: n=66; UNK: n=44) within FOCUS clinical trial cohort (**** $p < 0.0001$, t test). **B** Scatterplot showing correlation between desmoplastic stroma percentage determined from H&E and Fibroblast score determined by MCP-counter within the FOCUS clinical trial cohort (Spearman's rho = 0.64, $p < 2.2e-16$) coloured by CMS calls (CMS1: n=62; CMS2: n=155; CMS3: n=29; CMS4: n=66; UNK: n=44). **C** GSEA of Hallmark gene sets comparing samples with above median desmoplastic stroma percentage to samples with below median desmoplastic stroma percentage. Orange bars indicate gene sets significantly enriched in samples with above median desmoplastic stroma percentage while blue bars indicate gene sets significantly enriched in samples with below median desmoplastic stroma percentage (adjusted p-value < 0.02). Only the gene sets significantly and concordantly enriched in stroma or epithelium in both the LCM discovery and LCM validation cohorts are shown and gene sets are arranged in same order as in the LCM discovery cohort (Supplementary Figure 2A).

#### Supplementary Figure 4

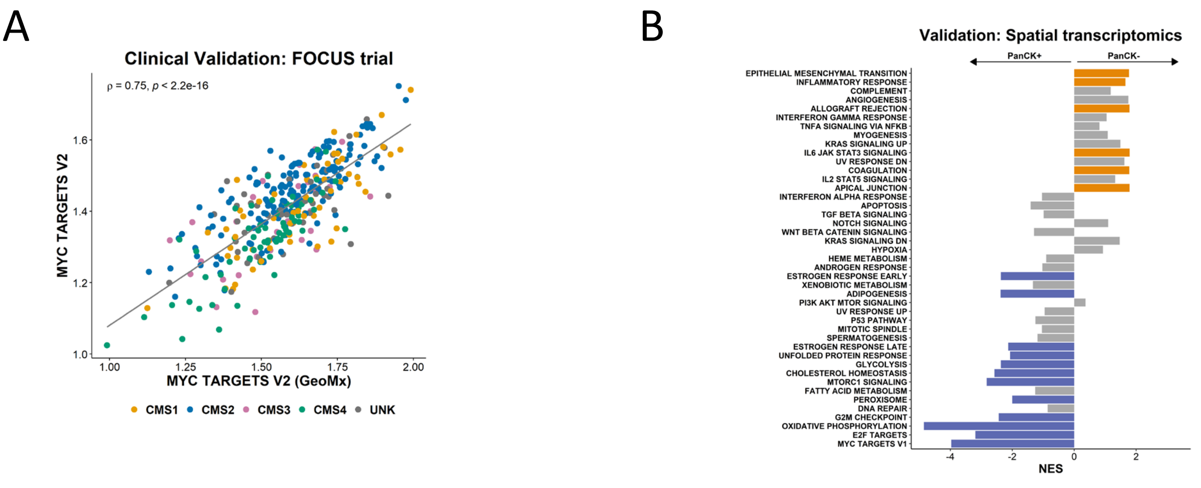


##### Supplementary Figure 4

**A** Scatterplot showing the correlation between ssGSEA scores for the full Hallmark MYC Targets V2 gene set and the GeoMx MYC Targets V2 gene set in the FOCUS clinical trial cohort (Spearman's rho = 0.75, $p < 2.2e\text{-}16$). Samples coloured by CMS subtype (CMS1: n=62; CMS2: n=155; CMS3: n=29; CMS4: n=66; UNK: n=44). **B** GSEA comparing PanCK- areas (stroma) to PanCK+ areas (epithelium) for the Hallmark gene sets (GeoMx versions). Orange bars indicate gene sets significantly enriched in PanCK- areas (stroma) (n=11) while blue bars indicate gene sets significantly enriched in PanCK+ areas (epithelium) (n=8) (adjusted p-value < 0.02).
